# Circadian-related Dynamics of the Endocannabinoid System in Male Mouse Brain

**DOI:** 10.1101/2025.04.15.648929

**Authors:** Dorit Farfara, Gil Moshe Lewitus, Ben Korin, Haitham Hajjo, Carni Lipson Feder, Liron Sulimani, Paula Berman, Shiri Procaccia, Hilla Azulay-Debby, David Meiri

**Author notes:** Corresponding author: Dorit Farfara, Department of Neurobiology and Immunology, Ruth and Bruce Rappaport Faculty of Medicine, Technion- Israel Institute of Technology, Haifa 31096, Israel.

## Abstract

Endocannabinoids (eCBs) and related lipids play crucial roles in brain function, including the regulation of circadian rhythms and sleep. To comprehensively map these molecules, we employed liquid chromatography high-resolution tandem mass spectrometry (LC/HRMS/MS) to quantify 78 lipids across 14 families in seven brain areas of male mice at four time points throughout the day (every six hours), and during sleep initiation. We found that most eCBs from the fatty acids (FAs) family, particularly arachidonic acid (AA), were highly abundant in the mouse brain in all brain areas and during the circadian rhythm. High eCBs abundance was shown in deeper brain areas, while temporal differences using the Cosinor analysis revealed 26 eCBs behaving in a circadian rhythm response, with linolenic acid (LnA) being the only lipid to show rhythmicity across all brain areas. Sleep initiation (ZT1) was associated with increased N-acylphosphatidylethanolamine phospholipase D (NAPE-PLD) activity and N-acylethanolamide (NAE) levels in the cortex and hippocampus, while wake extension (WEx) altered 2-monoacylglycerol (2-MAG) metabolism and increased cannabinoid receptor 1 (CB1) expression. These findings provide a detailed lipidomic map of eCBs and related lipids in the male mouse brain, highlighting their area-specific distribution, circadian regulation, and involvement in sleep/wake transitions. Given the link between sleep disruption and neurodegeneration, future studies should investigate whether the observed eCB dysregulation contributes to sleep disturbances in these conditions, and if targeting these pathways offers novel therapeutic strategies.

**Significant Statement:** This comprehensive study provides a high-dimensional map of eCBs and related lipids in the male mouse brain, revealing intricate spatial and temporal dynamics highlighting the role of these lipids in regulating fundamental physiological processes such as circadian rhythm and sleep.

## Introduction

The endocannabinoid (eCB) system is a ubiquitous endogenous neuromodulator signaling system with widespread functions in the brain and throughout the body^1,2^. The eCB system involves bioactive lipids known as endogenous cannabinoids, lipophilic signaling molecules, metabolic enzymes, and their receptors^3^. This diverse system is highly conserved^4^ and regulates many physiological systems, including circadian rhythmicity^5–7^, and sleep^8–10^. Distinct brain areas regulate these processes, synchronized by circadian rhythms^11,12^.

Most studies have focused on the two first-discovered and most familiar components of the classical eCBs: Arachidonoylethanolamide (AEA or anandamide)^13^, and 2-Arachidonoylglycerol (2-AG)^14,15^, and their cannabinoid receptors (CB1 and CB2)^16,17^. Today, additional eCBs and eCB-related lipids, receptors, and turnover enzymes, are considered a part of the extended eCB system ^18–21^. It is clear that the production of different eCBs varies between brain areas and is affected by physiological and cognitive experiences^22–24^. Furthermore, previous studies suggest that eCBs-mediated signaling varies with the time of the day^12^ or due to sleep deprivation^10,25^. AEA and 2-AG fluctuate throughout the day in phase with circadian rhythmicity^5,7,17,25^, as does the activity of synthesizing and hydrolyzing enzymes^26–28^. Furthermore, the density of the CB1 receptor was found to change with correlation to the time of day^29,30^, and sleep was altered in mice in which CB1 was knocked out^31^.

Here, we present a comprehensive endocannabinoidomics^32^ data set generated by advanced analytical techniques^21^. We conducted a large-scale screening to characterize changes in 78 eCBs, eCB-related lipids, and related compounds, in a circadian cycle and during sleep initiation. These eCB levels were analyzed across seven brain areas in naïve adult male mice in a six-hour interval over a 24-hour circadian cycle. We further assessed the levels of these eCBs and eCB-related lipids, metabolic enzymes, and their receptor, in brain areas during the first hour of sleep (sleep onset), and following a one-hour wake extension (WEx).

This comprehensive characterization of multiple eCBs, eCB-related lipids, and their metabolic pathways in the male mouse brain provides a valuable dataset that enhances our understanding of basic mechanisms involving circadian rhythms and sleep regulation, thus potentially opening new avenues in brain disorders involving sleep disturbances.

## Methods

### Mice

All experiments were performed under the National Institute of Health Guide for the Care and Use of Laboratory Animals. The Technion Administrative Panel approved all procedures and protocols satisfied Laboratory Animal Care. Adult male C57Bl/6 mice (https://www.inotivco.com/model/c57bl-6jola8-10 weeks of age; 20–25 gr) were maintained under Specific-Pathogen-Free (SPF) conditions. Females were not included in this study due to the fluctuations in endocannabinoid levels associated with the hormonal cycle^33,34^. Four mice were housed in each cage and held under controlled lighting (12:12 light-dark cycle, lights on at 09:00 AM), temperature (24±1°C), and humidity (30%–70%) conditions. Food and water were available ad libitum throughout the experiments. Throughout the manuscript, the light cycle refers to the mice sleeping phase (ZT—ZT6) and the dark cycle refers to their active phase (ZT12-ZT18). ZT0 is when lights turn on in the animal facility (i.e., the start of the light cycle, 9:00 AM), just before entering the mice sleep phase. ZT6 is six hours into the light cycle, during the mice sleep phase (3:00 PM). ZT12 is the beginning of the dark cycle when mice start their active phase (9:00 PM). ZT18 is after six hours into the dark cycle when the mice are still active (3:00 AM). The light-to-dark transition phase refers to ZT6-ZT12, and the dark-to-light transition phase refers to ZT18-ZT0

### ZT1 and Wake extension (WEx)

ZT1 indicates one hour after lights turn on, into the sleeping phase (initiation of the sleeping phase). WEx indicates one hour of gentle handling and inserting marbles into the cage to prevent sleep initiation while lights are on for an hour (sleep initiation disturbance).

### Tissue extraction and processing

At the end of each experimental time point, mice were euthanized by decapitation, and their brains were carefully extracted and placed immediately in ice-cold PBS. Seven brain areas (brainstem, cerebellum, cortex, hippocampus, striatum, thalamus, and hypothalamus) were dissected on a cooled aluminum tray under a surgical microscope, placed in a pre-weighed Eppendorf tube, and immediately snapped-frozen in liquid nitrogen. Tissue samples were kept in a −80°_C_ freezer until used for lipid extraction.

### Endocannabinoid quantification using LC/HRMS/MS

A minimal detection level was determined as 1 ng/gr per tissue. Standards were used as previously reported^21^.

### Endocannabinoid receptors and enzymes quantification using Real-Time PCR

RNA was extracted by cutting approximately 30 mg tissue sample and lysing in 1 ml TRI Reagent® (MRC Inc. TR 118). The lysate was added with 200 µl of chloroform followed by 15 min 12K RPM centrifugation. Approx. 500 µl of the upper aqueous phase was transferred to a clean tube to which 500 µl of 100% ethanol was added. The sample was loaded onto a Direct-zol column (Direct-zol™ RNA Miniprep kit, ZYMO RESEARCH) and purified according to the manufacturer’s instructions (including the optional DNAse step). Samples were run on a gel to verify no degradation had occurred and quantified on a Qubit 4 Fluorometer using Qubit™ RNA HS Assay Kit (Thermo-fisher Q32852). cDNA prepared with qScript® cDNA Synthesis Kit (Quantabio # 95047) using 1 µg of RNA in 20 µl reaction volume according to manufacturer protocol.

qPCR reactions were performed on a BIORAD-CFX384 using HOT FIREPol® EvaGreen® qPCR Mix Plus (Solis BioDyne, #08-24-00001) in a 10 µl reaction volume. Best universal reference genes were selected by extracting data from the mouse expression atlas for all tissues and selecting genes with mid-range expression in all tissues and the lowest calculated Coefficient of Variation values (mACTb, mAtp5f1, mEif2b1, mGusb, mHprt, mRps18, mSdha). A geometric mean value was calculated from all reference genes and used as a common normalization for all tissue samples. The experiment was conducted twice on at least three separate biological repeats to verify repetitive results.

### Experimental design and statistical analysis

Data were analyzed with GraphPad PRISM version 10 and are reported as mean ± standard error of the mean (s.e.m). When comparing compound levels across brain areas, we used one-way ANOVA, post hoc – Bonferroni, or ranks non-parametric assumption, with the Kruskal-Wallis (KW) rank sum test followed by Dunn’s test if the normalization test (Shapiro-Wilk test) was not significant (at least one parameter). Outliers were removed if ROUT<1%. To compare metabolites or areas within each family, we divided each value by the sum average of the total values of each metabolite or area (normalization), respectively. Only significant statistical results are presented. p ≤0.05 was considered statistically significant. * p<0.05, ** p<0.01, *** p<0.001, **\*\*\*\*** p<0.0001.

## Results

### eCBs and eCB-related lipid expression in brain areas at ZT0

To generate a comprehensive, high-dimensional map of eCBs and eCB-related lipids in the male mouse brain, we employed a liquid chromatography high-resolution tandem mass spectrometry (LC/HRMS/MS) analytical approach, as previously described^21^. Using this method, we evaluated the expression of 78 molecules across 14 lipid families (**Table 1**). Due to variations in synthesis and degradation pathways, the structure of eCBs can differ based on their polar hydrophilic head moiety (into families) or by the arachidonyl chain of the hydrophobic tail moiety (into names). Since the head moiety plays a key role in receptor binding, we categorized our analysis according to the hydrophilic head moieties within families.

**Table 1:**
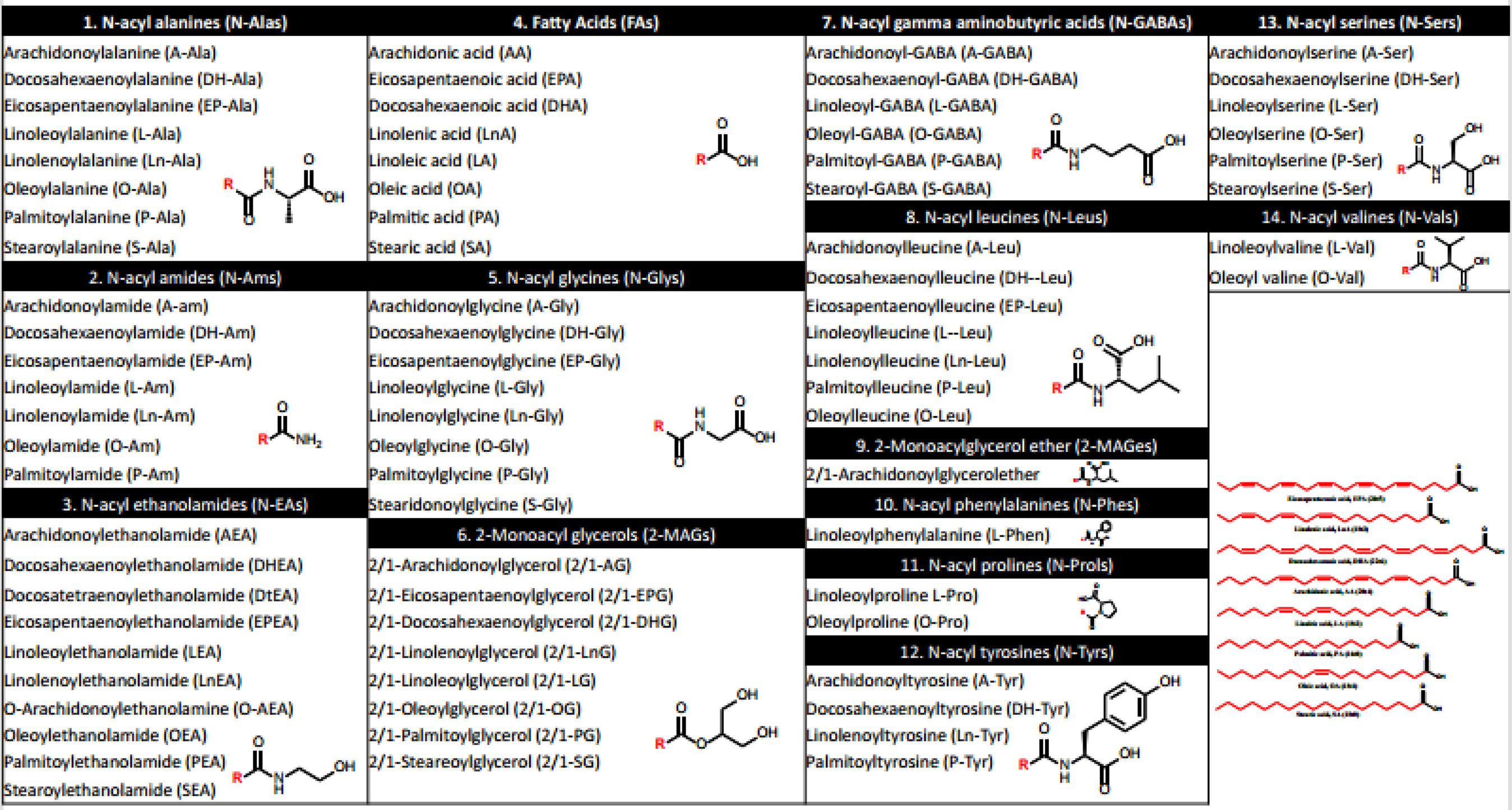
14 lipid families and their family members. The molecular structure is presented on the right. *R* represents the hydrophobic tail (names).

Our study focused on seven brain areas of the naïve male mice that are associated with circadian rhythms and sleep. We dissected seven brain areas: brainstem (BS), cerebellum (CB), and cortex (Cx), known as surface-related areas, and hippocampus (HC), striatum (ST), thalamus (TH), and hypothalamus (HT), known as deeper brain areas. We assessed the overall average concentrations of these molecules in weighed tissues (ng/g tissue) at light onset (Zeitgeber 0; ZT0) across these brain areas. Among 78 eCBs and eCB-related lipids, 43 from 10 different families were detected in more than two brain areas (**Figure 1A and Table 2**). Of these, 25 had available standards (marked with an asterisk in **Table 2**), enabling us to evaluate and compare their concentrations across different brain areas. The eCBs were categorized into three groups based on their concentration at ZT0: extremely high (>10^4), high (<10^4 and >10^2), and low (<10^2). The FA family members showed the highest expression compared to all analyzed eCBs and eCB-related lipids. Notably, Arachidonic acid (AA), a key structural component of many eCBs, showed the highest average expression across all brain areas. The extremely high concentrations of Docosahexaenoic acid (DHA), Palmitic-, Stearic-, and Oleic-acids (PA, SA, and OA) is consistent with previous reports ^35–38^. In accordance with existing literature^39^, high concentrations of Linoleic acid (LA), Eicosapentaenoic acid (EPA), and Linolenic acid (LnA) exhibited approximately 10-fold lower expression across all brain areas. The second most abundant lipid family in the brain was the 2-monoacylglycerols (2-MAGs) with 2/1-Eicosapentaenoylglycerol (2/1-EPG) and 2/1-Linoleoylglycerol (LG) exhibiting low concentrations in all brain areas. The N-Amides (N-Am) followed, with Linolenoylamide (Ln-A) and Arachidonoylamide (A-Am) showing low concentrations across all brain areas. All other lipid families, including members of the N-acylethanolamide (NAE) exhibited low expression in all brain areas consistent to previous results^40^.

**Figure 1.**
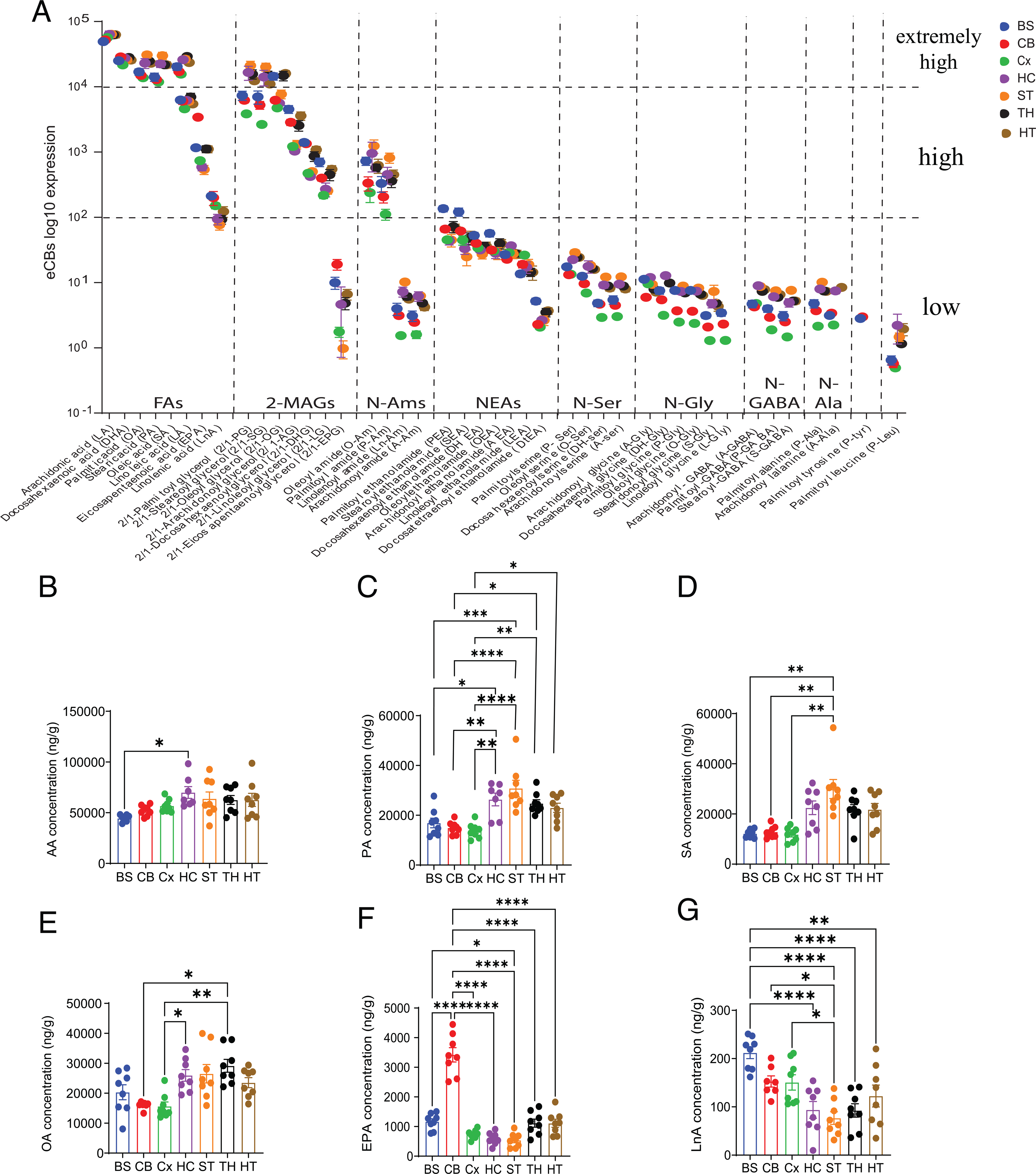
eCBs expression in ZT0 in all brain areas. **A.** log10 expression of eCBs families in brain areas at ZT0. Averaged values are presented in Table 2. **B.** AA concentrations (ng/g tissue) in ZT0 in all brain areas. (p=0.012 ANOVA, one way). **C.** PA concentrations (ng/g tissue) in ZT0 in all brain areas. (p<0.0001 ANOVA one way). **D.** SA concentrations (ng/g tissue) in ZT0 in all brain areas. (p<0.0001 ANOVA Kruskal-Walis). **E.** OA concentrations (ng/g tissue) in ZT0 in all brain areas. (p=0.0003 ANOVA Kruskal-Walis). **F.** EPA concentrations (ng/g tissue) in ZT0 in all brain areas. p<0.0001 (ANOVA, One way). G. LnA concentrations (ng/g tissue) in ZT0 in all brain areas. (p<0.0001 ANOVA one way).

**Table 2:** Descending average concentrations (ng/g tissue) at light onset (Zeitgeber 0; ZT0) of eCBs detected molecules in more than two brain regions. Asterisk (*) marks molecules with available standards.

| eCBs/brain areas | Brainstem | Cerebellum | Cortex | Hippocampus | Striatum | Thalamus | Hypothalamus |
| --- | --- | --- | --- | --- | --- | --- | --- |
| Arachidonic acid * | 49145.25 | 52218.13 | 56749.13 | 63387.39 | 63761.25 | 62540.75 | 62628.63 |
| Docosahexaenoic acid | 25072.25 | 28652.38 | 21593.50 | 27490.71 | 25792.88 | 28170.75 | 25710.75 |
| Oleic acid * | 20307.13 | 17053.25 | 15634.13 | 25943.13 | 26503.25 | 29140.38 | 23509.38 |
| Palmitic acid | 16886.25 | 14925.63 | 13765.38 | 23050.26 | 30792.13 | 24753.63 | 22994.63 |
| Stearic acid | 14139.25 | 12841.75 | 11822.38 | 22455.13 | 29928.00 | 21709.75 | 21763.38 |
| 2/1-Palmitoyl glycerol | 7360.18 | 6218.78 | 3841.32 | 16571.33 | 21143.10 | 15618.77 | 12380.31 |
| 2/1-Stearoyl glycerol | 7002.50 | 5231.38 | 2648.50 | 14089.71 | 20149.13 | 15404.38 | 11027.38 |
| 2/1-Oleoyl glycerol * | 14458.75 | 6226.88 | 4738.63 | 5593.25 | 7716.13 | 14790.63 | 16022.88 |
| Linoleic acid * | 6277.00 | 6022.88 | 4591.00 | 6199.29 | 5945.50 | 6985.25 | 5456.13 |
| 2/1-Arachidonoyl glycerol * | 4531.13 | 2851.50 | 1214.13 | 1031.50 | 1309.25 | 2558.25 | 3594.75 |
| Eicosapentaenoic acid | 1163.88 | 3420.00 | 736.13 | 587.63 | 538.00 | 1114.00 | 1110.38 |
| 2/1-Docosahexaenoyl glycerol | 1401.25 | 1341.25 | 471.75 | 423.00 | 502.13 | 862.25 | 1084.50 |
| N-Oleoyl amide * | 728.75 | 334.25 | 238.50 | 955.00 | 1242.25 | 573.25 | 624.20 |
| 2/1-Linoleoyl glycerol * | 703.25 | 398.63 | 217.00 | 270.00 | 254.00 | 450.63 | 545.00 |
| N-Palmitoyl amide | 330.70 | 205.63 | 111.29 | 452.63 | 817.86 | 359.71 | 451.00 |
| Linolenic acid * | 211.88 | 199.75 | 150.63 | 93.70 | 76.68 | 92.49 | 122.44 |
| Palmitoyl ethanolamide * | 135.74 | 66.03 | 45.19 | 43.86 | 44.84 | 71.94 | 67.99 |
| Stearoyl ethanolamide * | 120.90 | 61.59 | 44.60 | 33.10 | 24.83 | 47.60 | 49.75 |
| Docosahexaenoyl ethanolamide * | 52.76 | 40.03 | 34.01 | 31.66 | 27.04 | 36.01 | 36.20 |
| Oleoyl ethanolamide * | 56.66 | 31.51 | 30.41 | 28.20 | 28.41 | 39.70 | 38.98 |
| Arachidonoyl ethanolamide * | 26.91 | 22.78 | 28.61 | 36.56 | 27.39 | 28.04 | 27.35 |
| N-Palmitoyl serine | 17.41 | 13.19 | 13.41 | 22.50 | 28.79 | 24.36 | 24.34 |
| Linoleoyl ethanolamide * | 13.57 | 18.99 | 26.29 | 17.21 | 18.91 | 14.46 | 14.38 |
| N-Oleoyl serine | 12.49 | 9.60 | 6.92 | 17.71 | 18.34 | 16.40 | 15.91 |
| N-Arachidonoyl glycine * | 11.20 | 5.94 | 9.55 | 11.93 | 9.37 | 7.91 | 7.14 |
| N-Palmitoyl glycine * | 7.56 | 3.69 | 2.48 | 19.00 | 8.82 | 6.93 | 7.55 |
| 2/1-Eicosapentaenoyl glycerol | 9.96 | 19.00 | 1.75 | 4.63 | 0.98 | 4.84 | 6.61 |
| N-Docosahexaenoyl serine | 4.84 | 4.66 | 2.92 | 9.17 | 12.12 | 8.87 | 8.23 |
| N-Arachidonoyl serine * | 5.45 | 4.49 | 3.01 | 9.46 | 12.25 | 8.83 | 7.98 |
| N-Arachidonoyl GABA * | 4.70 | 4.24 | 4.77 | 8.95 | 8.71 | 7.33 | 7.82 |
| N-Docosahexaenoyl glycine * | 7.50 | 5.45 | 3.14 | 12.77 |  | 9.84 |  |
| N-Palmitoyl alanine | 4.77 | 3.70 | 2.14 | 7.98 | 10.11 | 7.42 | 7.32 |
| N-Oleoyl glycine * | 7.57 | 3.62 | 2.36 | 7.39 | 8.24 | 6.63 | 6.34 |
| N-Linolenoyl amide * | 3.97 | 3.12 | 1.54 | 7.23 | 10.19 | 6.10 | 5.57 |
| N-Palmitoyl GABA | 3.97 | 2.93 | 1.89 | 6.05 | 7.49 | 5.95 | 6.54 |
| N-Arachidonoyl alanine * | 3.14 | 3.39 | 2.23 | 7.45 |  |  | 8.43 |
| N-arachidonoyl amide * | 3.10 | 2.46 | 1.60 | 6.18 | 6.25 | 4.85 | 4.20 |
| N-Stearoyl GABA | 3.09 | 2.50 | 1.47 | 4.92 | 7.59 | 5.21 | 5.22 |
| N-Stearidonoyl glycine * | 3.13 | 2.08 | 1.30 | 4.77 | 7.29 | 4.84 | 4.32 |
| Docosatetraenoyl ethanolamide * | 5.19 | 2.28 | 2.07 | 2.65 | 2.64 | 3.49 | 3.68 |
| N-Palmitoyl tyrosine | 2.81 | 2.98 |  |  |  |  |  |
| N-Linoleoyl glycine | 3.44 | 2.31 | 1.30 |  |  |  |  |
| N-Palmitoyl leucine | 0.65 | 0.57 | 0.49 | 2.21 | 1.47 | 1.15 | 1.94 |

To analyze the abundance of eCB families in each brain area, we averaged and normalized their concentrations per tissue weight and found that deeper brain areas exhibited high eCB expression (**Supplementary Figure 1A**), consistent with previous findings^41,42^. When analyzing each eCB family separately, we found no significant differences in expression across brain areas within the FA family (**Supplementary Figure 1B**). However, 2-MAG and N-Am families showed significantly low eCB expression in the Cx (**Supplementary Figure 1C-D**), while the CB exhibited high eCB levels within the NAE family (**Supplementary Figure 1E**). The remaining lipid families (N-Gly, N-Ser, N-GABA) showed higher abundance in deeper brain areas than surface ones (**Supplementary Figure 1F-H**).

Upon further examination of specific changes in individual eCB levels across brain areas, we found that within the FA family, AA exhibited minimal variation between brain areas, with a trend towards higher concentrations in deeper parts of the brain, specifically the HC (**Figure 1B**). Similarly, PA, SA, and OA demonstrated significantly higher expression in deeper brain areas compared to surface-related areas (**Figure 1C-E**). In accordance to previous data indicating that DHA is highly expressed than EPA^39^, DHA did not show any significant changes within specific brain areas (data not shown), however, EPA showed a notable increase in the CB compared to other brain areas (**Figure 1F**). Additionally, LnA demonstrated higher expression in surface-related areas, specifically in the BS (**Figure 1G**), suggesting distinct functional roles of FA family members across different brain areas.

Individual eCBs within the 2-MAGs revealed that 2/1-Palmitoylglycerol (2/1-PG) and 2/1-Steareoylglycerol (2/1-SG), had significantly higher expression in deeper brain areas, whereas 2/1-Oleoylglycerol (2/1-OG), 2/1-Arachidonylglycerol (2/1-AG), 2/1-Docohexanoylglycerol (2/1-DHG) and 2/1-Linoleoylglycerol (2/1-LG) showed high expression in the BS, TH, and HT compared to Cx, HC, and ST. 2/1-Eicosapentaenoilglycerol (2/1-EPG) exhibited higher expression in BS and CB (**Supplementary Figure 2A**), following a similar pattern to that observed with EPA. Within the N-Am family, highly expressed Oleylamide (O-Am) and Palmitoylamide (P-Am) and low-expressed Linolenoylamide (Ln-Am) and Arachidonoylamide (A-Am) followed the same pattern, showing higher expression in deeper areas of the brain (**Supplementary Figure 2B**). Most of the 2-NAE family members showed the BS to be relatively higher in expression, except for AEA and Linoleoylethanolamide (LEA), which demonstrated a similar trend as AA (**Supplementary Figure 2C**). The rest of the lipid families (N-Ser, N-Gly, and N-GABA) followed the trend, showing higher expression in deeper areas of the brain (**Supplementary Figure 2D-F**).

### A circadian profile of eCBs and eCB-related lipid in brain areas

Based on previous reports, we tested the fluctuation, range, and patterns of changes during a circadian rhythm cycle^5,6,43^. We examined variations in eCB concentrations at 4 time points across the circadian cycle, spanning both the light (ZT0-ZT6, sleep phase) and dark (ZT12-ZT18, active phase) periods (**Scheme 1**). Total normalized eCBs showed significant changes in the light-to-dark (ZT6-ZT12) and dark-to-light phase transitions (ZT18-ZT0) in the CB, and ST (**Figure 2A-B**). The Cx showed significant changes in the light phase transition and within the dark phase transition changes were detected in the BS and TH (**Supplementary Figure 3A, B, and D**). Furthermore, eCBs in the Cx, ST, BS, TH, and HT were affected by a typical 12-hour light/dark (ZT0-12, and ZT6-18) circadian cycle **(Supplementary Figure 3).** Interestingly, the total averaged normalized eCB levels in the HC did not show any significant changes throughout the circadian cycle (**Supplementary Figure 3C**). By evaluating changes in averaged family members during the circadian cycle across specific brain areas, we observed a progressive increase in the FA family members (triangle-dotted columns) at ZT18 (**Supplementary Figure 4**), however, no changes were detected in the TH (data not shown). 2-MAGs (circle-dotted columns) decreased gradually with the progression of the circadian cycle with no changes in the ST, whereas N-Ams (starred-dotted columns) and NAEs (square-dotted columns) increased progressively throughout the circadian cycle in all brain areas.

**Scheme 1.**
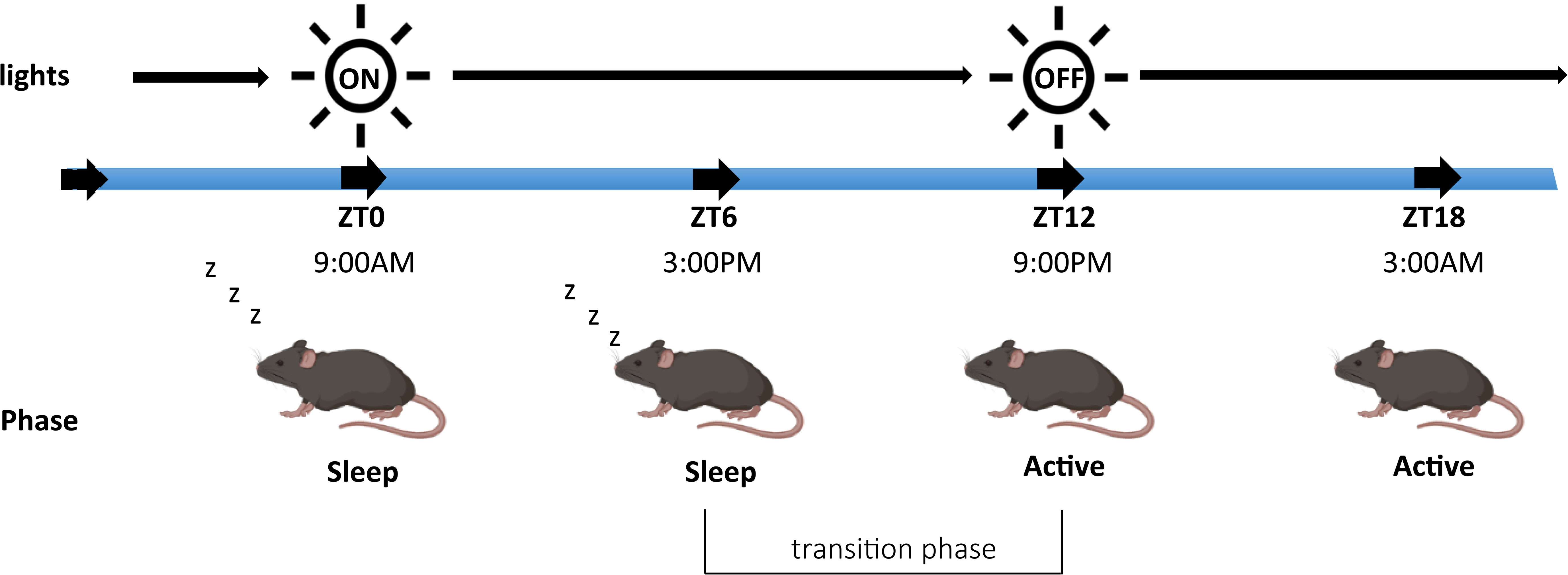
Circadian cycle diagram.

**Figure 2:**
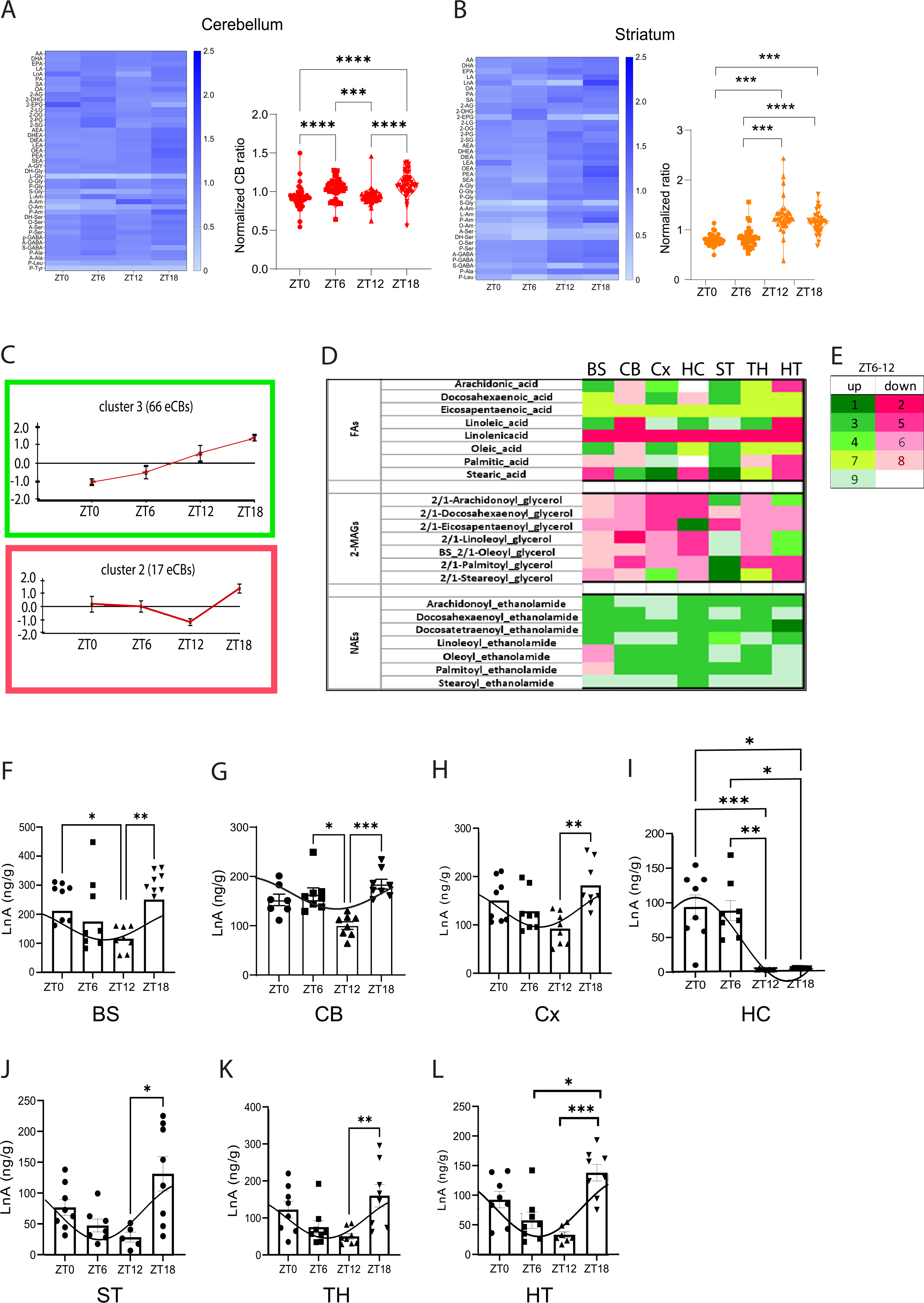
eCBs expression in circadian rhythm in brain areas. A-C. Left: Heat maps of eCBs from the CB, CT, and ST. **A-C.** Right: Quantification of eCBs during the circadian cycle in the eCBs from the CB (p<0.0001, ANOVA, Kruskal-Wallis test), CT (p<0.0001, ANOVA, Kruskal-Wallis test), and ST (p<0.0001, ANOVA, Kruskal-Wallis test). **C.** Clusters 3 and 2 show 66 and 17 eCBs within brain areas demonstrating progressive elevation (green frame) and decreased (red frame) expression patterns. **D.** FAs, 2-MAG, and NAEs expression in the transition phase (ZT6 to ZT12) demonstrating a decrease (red) or increased (green) concentration levels. **F.** Numbers 1-9 indicate the cluster types (together with Figure 2-3). **G-M.** Cosinor plots showing LnA expression in circadian fluctuation in BS (p=0.0009 ANOVA one-way), CB (p=0.0003 ANOVA Kruskal-Wallis test), CT (p=0.0021 ANOVA one-way), HC (p<0.0001 ANOVA Kruskal-Wallis test), ST (p=0.0044 ANOVA one-way), TH (p<0.0001 ANOVA one-way), HT (p=0.0045 ANOVA Kruskal-Wallis test).

Using k-means clustering, we grouped eCBs into nine clusters with high homogeneity based on their dynamic in the circadian pattern (similarity>0.913) and differentiated them based on their expression patterns during the light-to-dark (sleep-to-active) transition phase in a specific brain area (increased – green, decreased – red) (**Figure 2C-D and Supplementary Figure 5A-B**). The majority of the eCBs (66 eCBs) followed the same pattern as cluster 3, showing consistent elevation throughout the circadian cycle (**Figure 2C**). However, 17 eCBs followed the pattern of cluster 2, showing changes in the pattern expression between light-to-dark (ZT6-to-ZT12) (**Figure 2C**), 21 eCBs followed cluster 1, and 27 followed cluster 8 (**Supplementary Figure 5A-B).** Among them, LnA exhibited a decreased expression across all brain areas during the light-to-dark transition phase, while EPA followed cluster 7 (**Figure 2D-E**). We further found that 96% of all eCBs in the NAE family increased across all brain areas during the light-to-dark transition phase, while 78% of 2-MAG family members showed a decrease in ZT6-ZT12.

A Cosinor test determined which of the specific eCBs corresponded to a sine curve, indicating significant circadian fluctuations as previously suggested^44^. Out of 78 eCBs, 26 passed the cosinor test, with the majority being present in the Cx and TH and were mostly members of the NAEs family (**Supplementary Figure 6**). Interestingly, the only member of the FA family that passed the cosinor test in all brain areas was the LnA (**Figure 2F-L**), showing a dramatic decrease within the transition phase (ZT6-ZT12) specifically in the HC (**Figure 2I**), indicating a possible role in circadian regulation.

### Metabolic regulation of eCBs in sleep initiation and wake extension

To identify the metabolomic effects in the eCB system caused by light transition, we analyzed the gene expression of related enzymes and receptors after light transition, entering the sleep phase (ZT0), after an hour of light into the sleep phase (ZT1), and when the initiation of sleep was deliberately deprived for an hour (WEx) while the lights are on (**Scheme 2**). We conducted this experiment based on previous data showing this does not induce stress^45^. Specifically, we examined the expression of key metabolizing enzymes (biosynthesis and degradation) and receptors known to play a role in eCB pathways. *N-Acylphosphatidylethanolamine phospholipase D (NAPE-PLD)* synthesis enzyme, catalyzes the precursor N-acyl-phosphatidylethanolamines (NAPEs) to synthesize 2-NAEs^46^ (**Figure 3A**). NAPE-PLD was significantly increased during sleep initiation in the Cx and HC (**Figure 3B-C**). As a result, levels of 2-NAEs in the Cx and HC were increased within one hour of sleep (**Figure 3D-E and Supplementary Figure 7A**), while disruption of sleep did not synthesize eCBs, suggesting 2-NAE is regulated during sleep initiation in the Cx and HC. In contrast, within the degradation pathway (**Figure 3F**), 2/1-AG, 2/1-DHG, and 2/1-EPG were decreased in the CB in the first hour of sleep (ZT1) (**Figure 3G and Supplementary Figure 7B**), probably as a result of an increase in the *Monoacylglycerol lipase (MAGL)* degrading enzyme (**Figure 3H**), which hydrolysis 2-MAGs into FAs and glycerol^47^. However, 2/1-SG in the HT was increased in WEx compared to ZT0 (**Figure 3I**), suggesting decreased enzymatic activity of MAGL in WEx as seen in **Figure 3J**. The degrading enzyme of AEA, the *fatty acids amide hydrolase (FAAH)* was increased in ZT1 in the CB and BS compared to ZT0 (**Supplementary Figure 7C).** As for the receptors, cannabinoid receptor 1 (*CB1)* in the HC was increased during sleep initiation and WEx (**Figure 3K**), whereas cannabinoid receptor *2 (CB2)* in the ST was decreased in ZT1 (**Figure 3L**). *Peroxisome proliferator-activated receptors (PPARs)* regulating immune and cellular homeostasis^48^ increased significantly during sleep initiation in the BS (**Figure 3M**), and *G Protein-Coupled Receptor* (GPRs) known to be expressed in neurons^49^, were increased in ZT1 and WEx in the HC (**Figure 3N**). Receptors controlling heat regulation, such as *Transient Receptor Potential Cation Channel Subfamily V members (TRPVs*) and serotonergic receptors *Htr1A and Htr2A* changes are presented in **Supplementary Figure 7D**. Furthermore, by evaluating the changes in eCBs during ZT1 and WEx we revealed that most of the eCBs that were affected during the initiation of sleep (ZT1) were in the CB, whereas WEx influenced the eCBs in the deeper areas of the brain (**Supplementary Figure 8**).

**Scheme 1.**
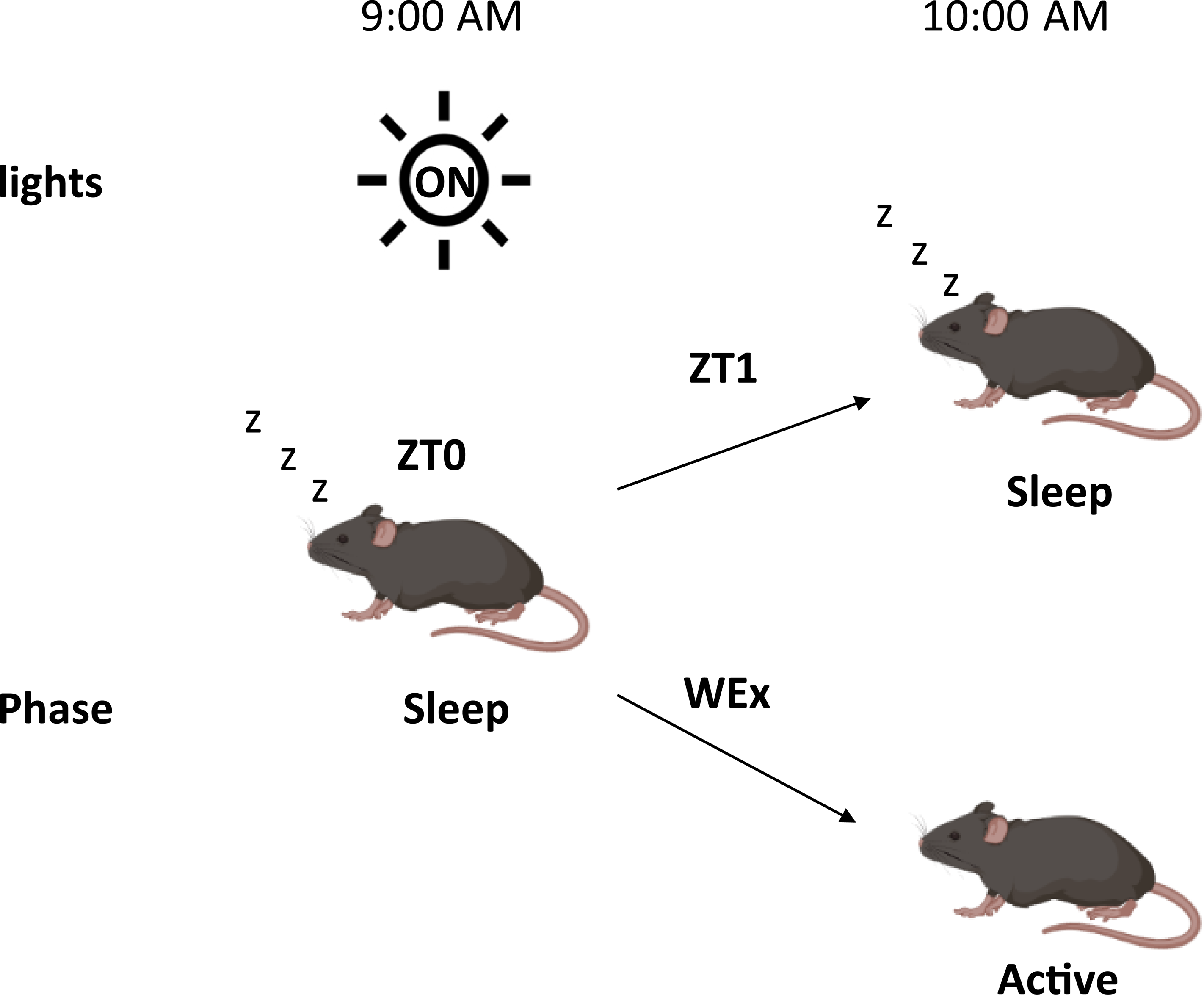
ZT0, ZT1, and WEx diagram.

**Figure 3:**
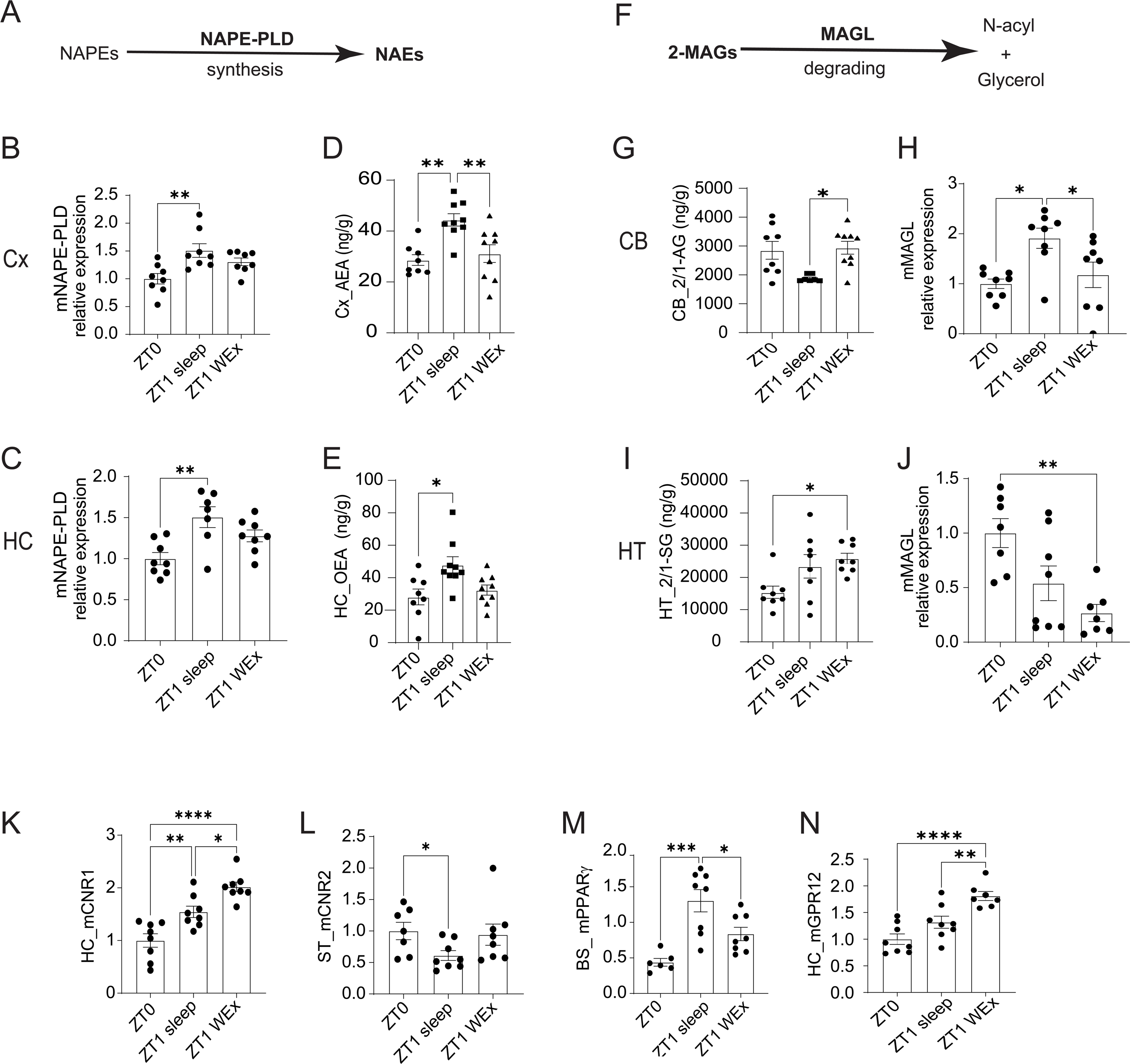
eCBs metabolic pathways during sleep initiation (ZT1) and wake extension (WEx) in the brain. **A.** Schematic representation of NAPE-PLD synthesis pathway. **B.** Gene expression of NAPE-PLD in the CT (p=0.005, ANOVA one way). **C.** Gene expression of NAPE-PLD in the HC (p=0.003, ANOVA one way). **D.** Levels of AEA in the CT (p=0.001, ANOVA one way). **E.** Levels of OEA in the HC (p=0.009, ANOVA one way). **F.** Schematic representation of MAGL degradation pathway. **G.** Levels of 2/1-AG in the CB (p=0.013, ANOVA Kruskal-Wallis). **H.** Gene expression of MAGL in the CB (p=0.008, ANOVA one way). **I.** Levels of 2/1-SG in the HT (p=0.02, ANOVA one way). **J.** Gene expression of MAGL in the HT (p=0.003, ANOVA one way). **K.** Gene expression of CNR1 in the HC (p<0.0001, ANOVA one way). **L.** Gene expression of CNR2 in the ST (p=0.036, ANOVA Kruskal-Wallis). **M.** Gene expression of PPARγ in the BS (p=0.0003, ANOVA one way). **N.** Gene expression of GPR12 in the HC (p<0.0001, ANOVA one way).

## Discussion

Most studies evaluating the physiological roles of endocannabinoids (eCBs) have primarily focused on anandamide (AEA) ^13^ and 2 AG^14^, the two earliest discovered eCBs. However, many other endogenous molecules, similar to eCBs in structure and metabolism, have been identified and included in the related eCB system. Characterizing these molecules across specific brain areas in circadian rhythm and sleep is crucial in understanding normal cycles and pathological outcomes, as previously demonstrated^40^. Using genomic and metabolomics tools^21^, we characterized the profiles and dynamics of 78 eCBs across seven brain areas in naïve male mice, sampled at four distinct time points throughout the day. This analysis also considered the effects of sleep onset and wake extension of the initial light change, providing insights into the temporal and area-specific regulation.

Our results demonstrate a comprehensive lipidomic profile of eCBs and related lipids in the male mouse brain, revealing intricate areaal distribution patterns and dynamic circadian fluctuations. Consistent with previous reports, we found that fatty acids (FAs) exhibited the highest overall expression, with arachidonic acid (AA) being the most abundant across all brain areas, particularly in the hippocampus (HC), reinforcing its key role in eCB metabolism^50^. Notably, while relatively low in concentration, eicosapentaenoic acid (EPA) and linolenic acid (LnA) showed exceptionally high levels in the cerebellum (CB) and brainstem (BS). This is significant given the importance of omega-3 polyunsaturated fatty acids, including DHA, LnA, and EPA, for brain function, development, and cognitive performance^51–54^ and brain disorders^55^.

The observed regional differences in eCB abundance, with deeper brain areas generally displaying higher levels, align with previous reports. However, NAEs were highly expressed in the CB, suggesting a specific role of these eCBs in cerebellar function. Interestingly, AEA and LEA did not follow the same pattern, highlighting the importance of studying non-canonical eCBs.

One challenge in circadian research is differentiating between fluctuations due to circadian control and other physiological processes, such as stress ^23^ or satiety^56^. We found that eCBs in the CB and striatum (ST) were dramatically elevated during light transition phase (ZT6-ZT12 and ZT18-ZT0), suggesting these brain areas are influenced by light/activity transition. Within the FA family, EPA and LnA showed a consistent increase and decrease across all brain areas within the light transition phase, indicating their involvement of these molecules in light transition. NAE members showed overall increased changes in the light transition phase, while 2-MAGs showed a general decrease, suggesting a specific family regulation depending on the timing of the day. Our observation supports the finding of Valenti et al.^25^, where AEA is higher during the dark period while 2-AG is higher during the light period.

Furthermore, Cosinor analysis revealed that only 26 of 78 analyzed eCBs exhibited significant circadian rhythmicity, primarily from the FA family in the cortex and thalamus. Notably, LnA was the only eCB to show rhythmicity across all brain areas, underscoring its potential role in circadian regulation. To discriminate changes that are due to light transition, circadian regulation, and sleep, we analyzed eCBs after the first hour of sleep (ZT1) and wake extension (WEx). Of note, a one-hour extension of the wake period is not considered to induce stress^45^. It is generally accepted that eCB signaling is regulated on-demand by their biosynthesis and catabolism within the plasma membrane from phospholipid precursors^57^. We observed changes in the mRNA levels of metabolic enzymes, with NAPE-PLD in the cortex and hippocampus affected by sleep, and MAGL affected by wake extension. Supporting previous findings, CB1 and CB2 receptors in the hippocampus and striatum were regulated by sleep^58,59^. Moreover, arachidonoyl-type lipids such as O-arachidonoyl-ethanolamine (O-AEA, virodhamine) were found to bind to the classical cannabinoid receptors CB1 and CB2^60^ whereas AEAs, 2-AGs, and their analogs were shown to bind also non-classical cannabinoid receptors, including other GPCRs^3,61–63^, PPARs^64^, TRPs^65^, serotonergic receptors^66^. We found that most receptors in the HC were affected by wake extension, suggesting that sleep disruption, rather than light transition, influences eCB signaling in this area.

To conclude, as the amounts of eCBs change rapidly with low absolute thresholds, our results should be taken with some methodological reservations. Technical procedures and postmortem effects might have affected eCB synthesis and degradation. Also, the commonly used C57Bl/6 mice, known for impaired melatonin biosynthesis, might have influenced our findings^53^.

In summary, this study provides a detailed dataset of eCBs and related lipids, revealing their spatial and temporal distribution, in circadian regulation, and involvement in sleep/wake transitions. Our findings suggest that these lipids play a crucial role in regulating brain function and behavior and provide a foundation for future research aimed at elucidating the precise mechanisms by which they exert their effects. Considering the established link between sleep disruption and neurodegeneration, our findings highlight the need to explore whether the observed endocannabinoid dysregulation plays a causative role in the sleep disturbances that manifest in these debilitating conditions.

## Acknowledgments

We thank Prof. Asya Rolls for all the support. All authors made a direct and meaningful contribution to the paper.

## Author contributions

Conceptualization: DF, GML, HAD, DM

Methodology: DF, GML, BK, HH, PB

Investigation: DF, CL, LS, DM

Visualization: DF, GML, SP

Supervision: DM

Writing – original draft: DF, GML, SP, DM

Writing – review & amp; editing: DF, GML, SP, DM

