## Supplementary figures and images for "Circadian-related Dynamics of the Endocannabinoid System in Male Mouse Brain"

### Supplementry figure1

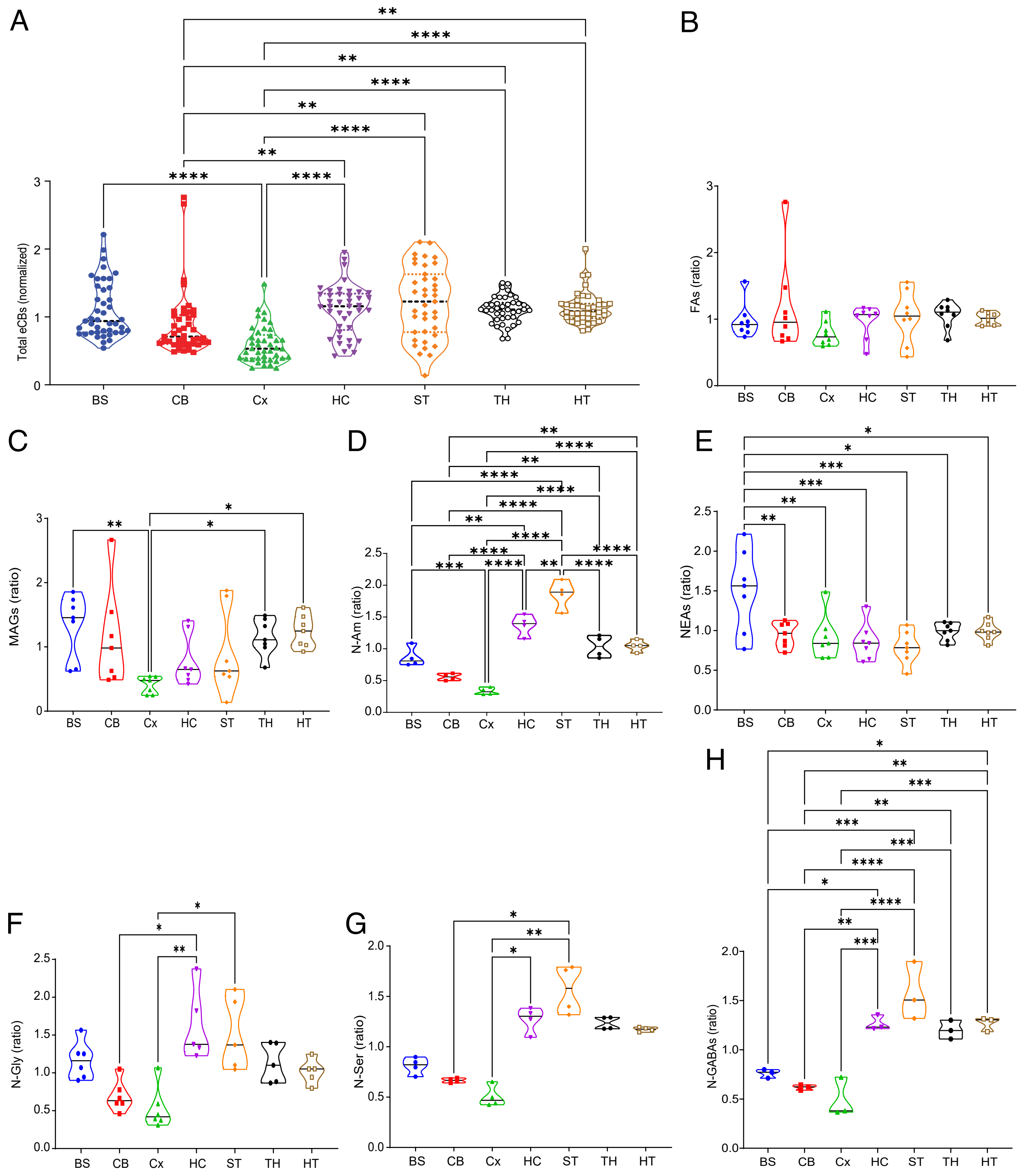

### Supplementry figure 2

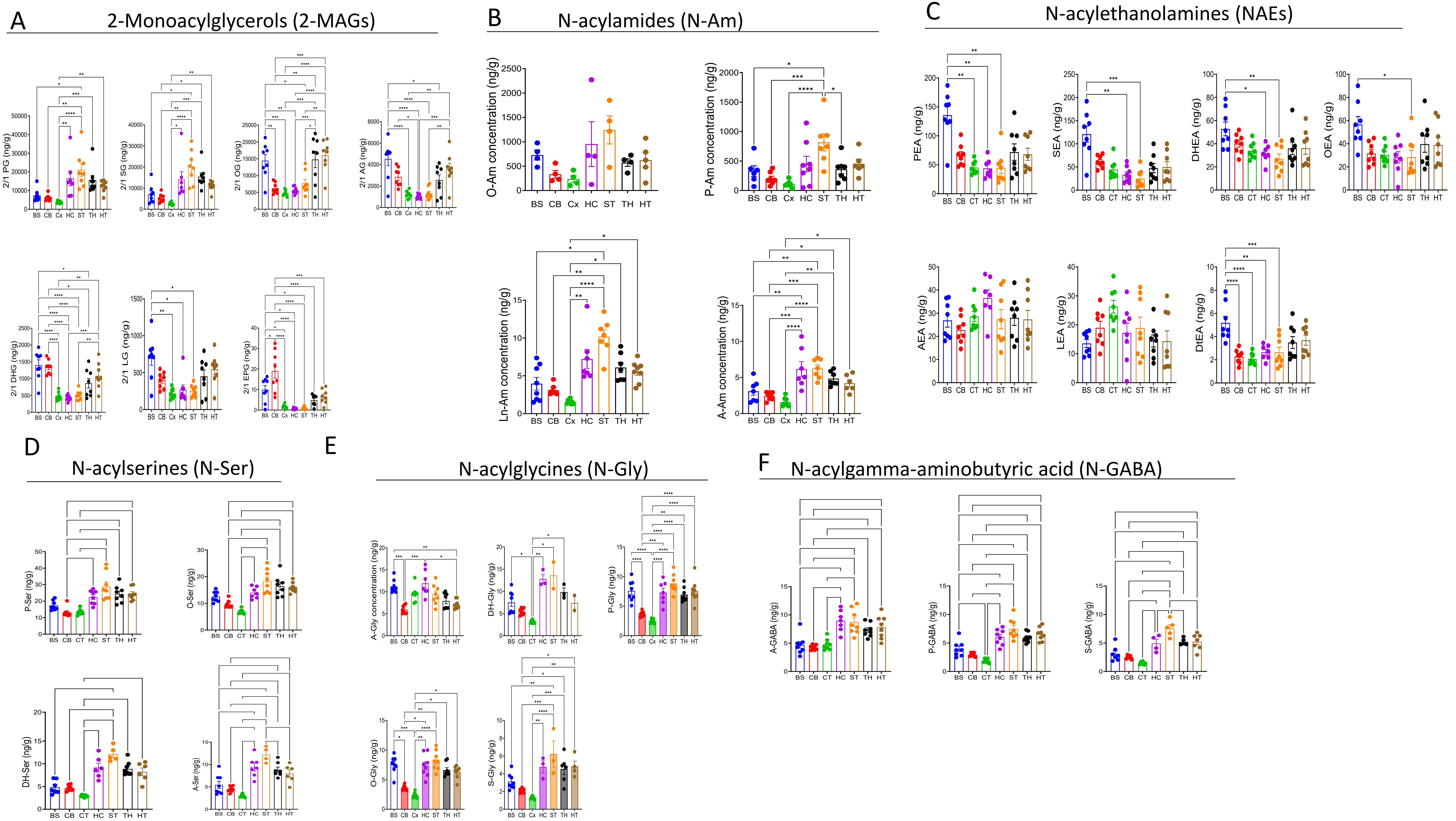

### Supplementry figure 3

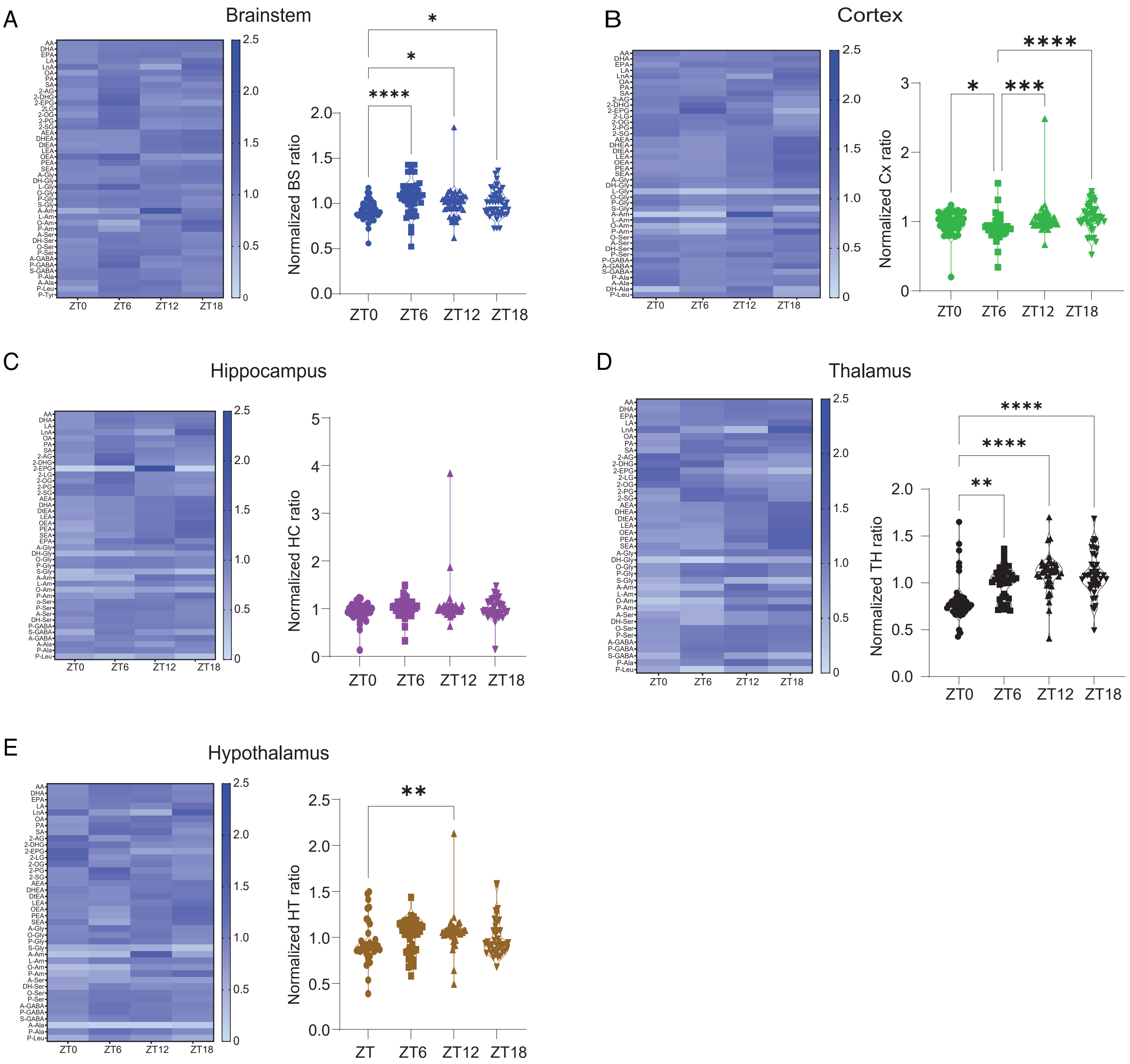

### Supplementry figure 4

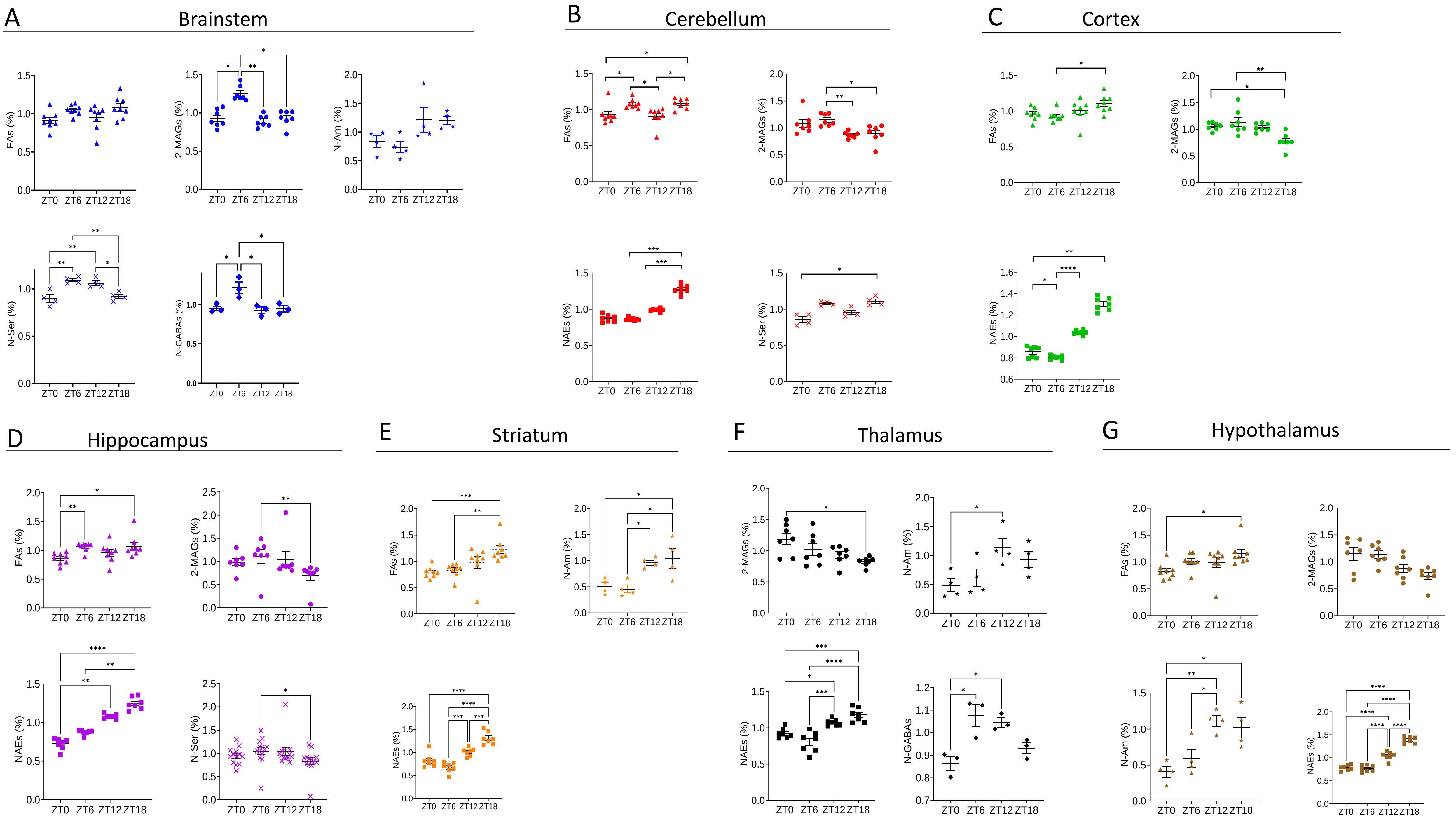

### Supplementry figure 6

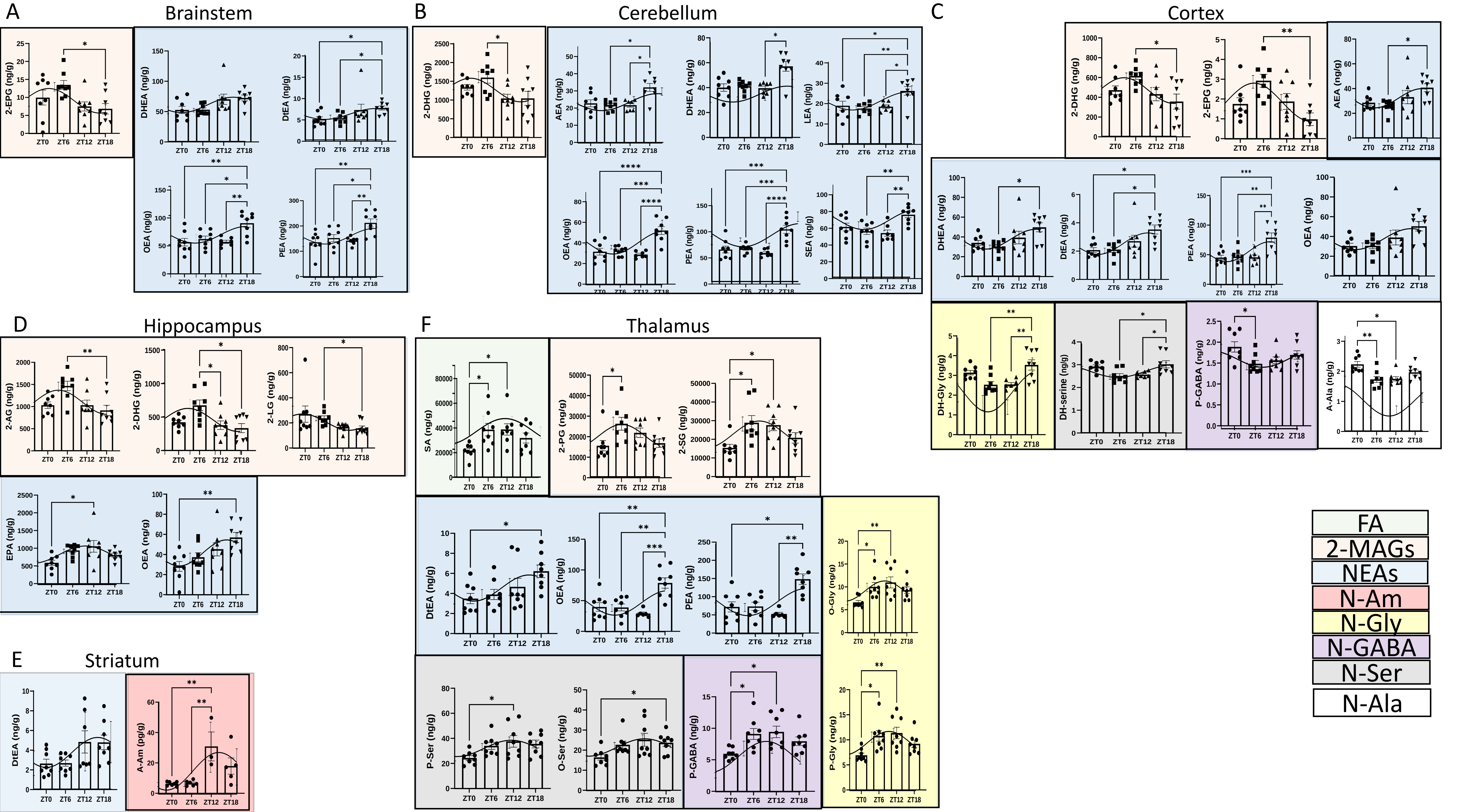

### Supplementry figure 7

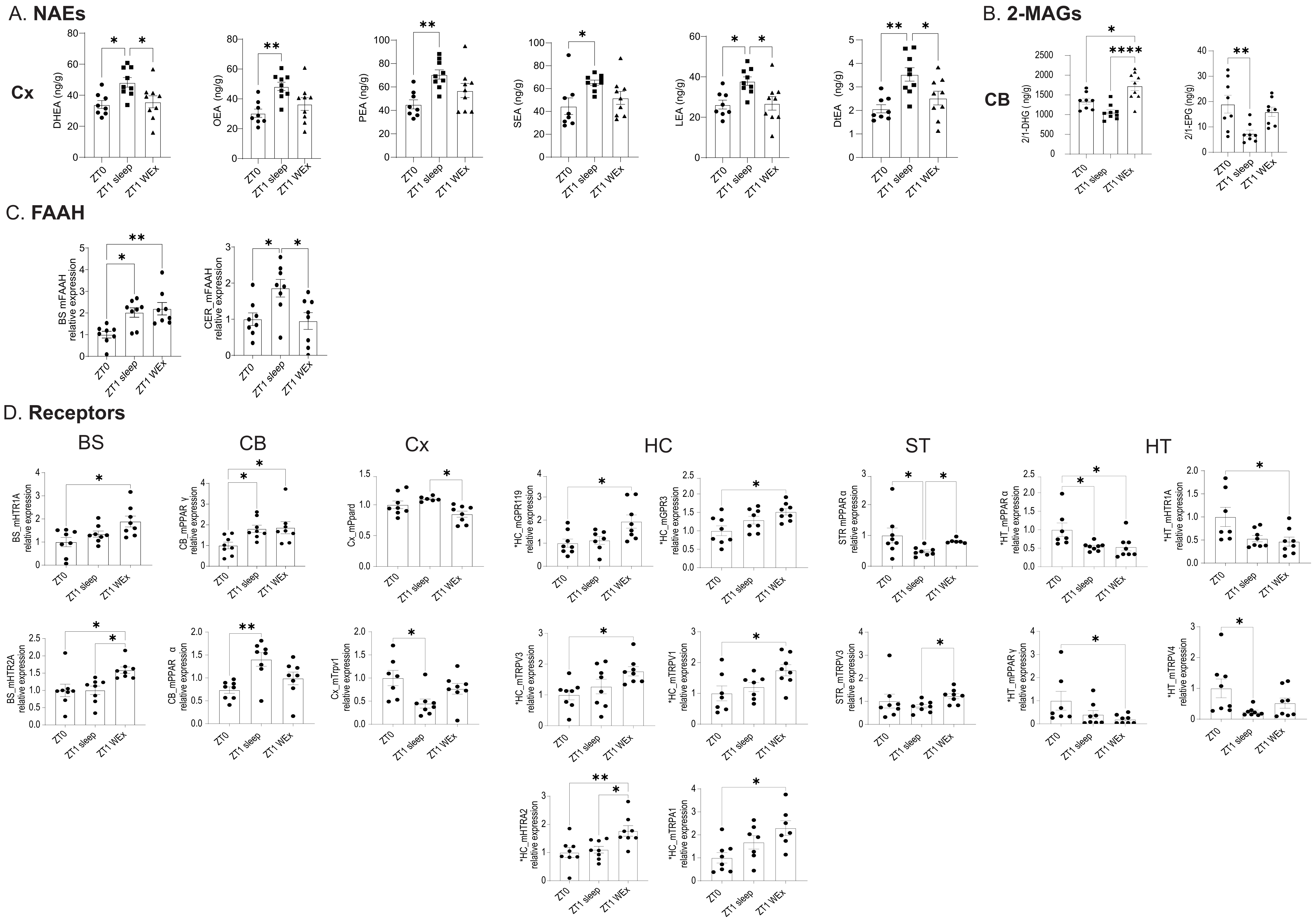

### Supplementry figure 8

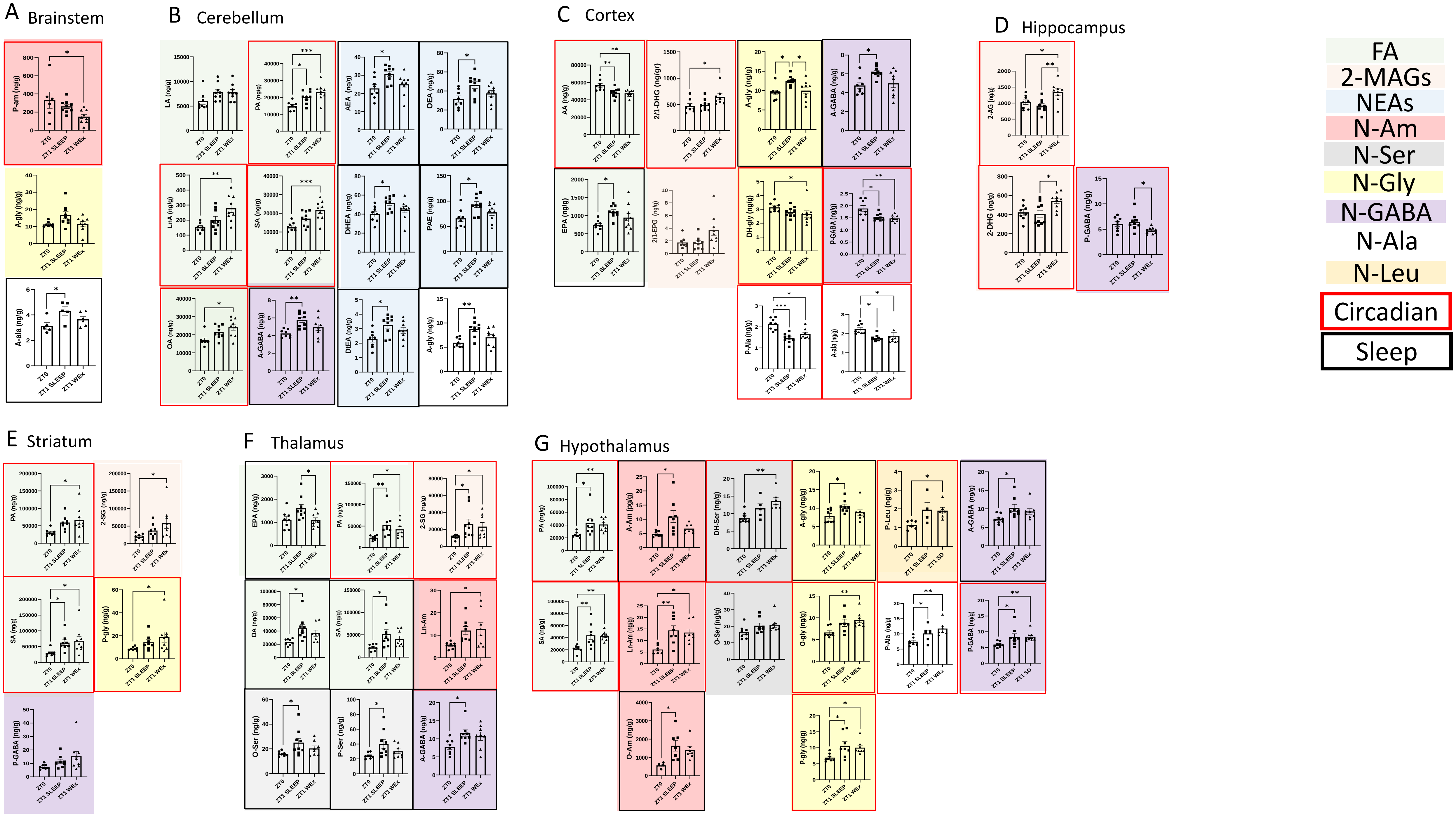
