## Supplementry figure 5 for "Circadian-related Dynamics of the Endocannabinoid System in Male Mouse Brain"

A

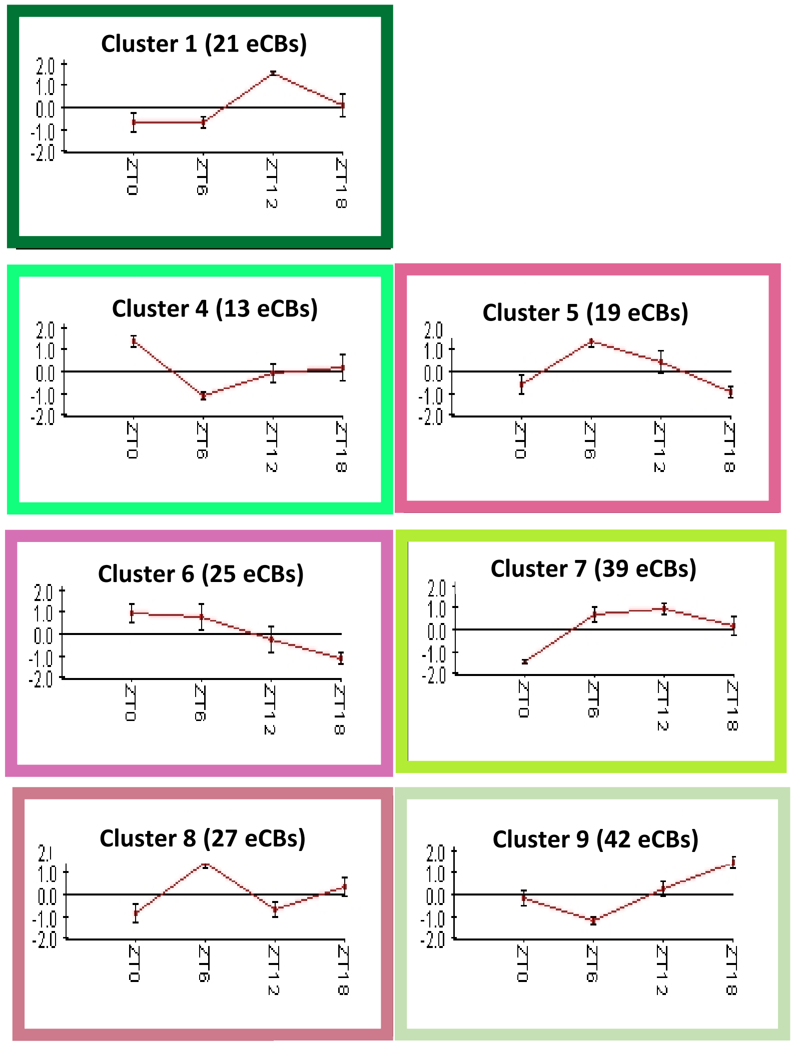


B

**Clustering Info:**
Algorithm: K-Means
Number of clusters (K): 9
Overall Average Homogeneity: 0.913
Overall Average Separation: 0.011
Number of clusters: 9
Number of singletons: 0

| ID | Name | Size | Homogeneity |
|---|---|---|---|
| 1 | Cluster_1 | 21 | 0.904 |
| 2 | Cluster_2 | 17 | 0.879 |
| 3 | Cluster_3 | 66 | 0.939 |
| 4 | Cluster_4 | 13 | 0.884 |
| 5 | Cluster_5 | 39 | 0.905 |
| 6 | Cluster_6 | 25 | 0.837 |
| 7 | Cluster_7 | 39 | 0.929 |
| 8 | Cluster_8 | 27 | 0.909 |
| 9 | Cluster_9 | 42 | 0.939 |

C

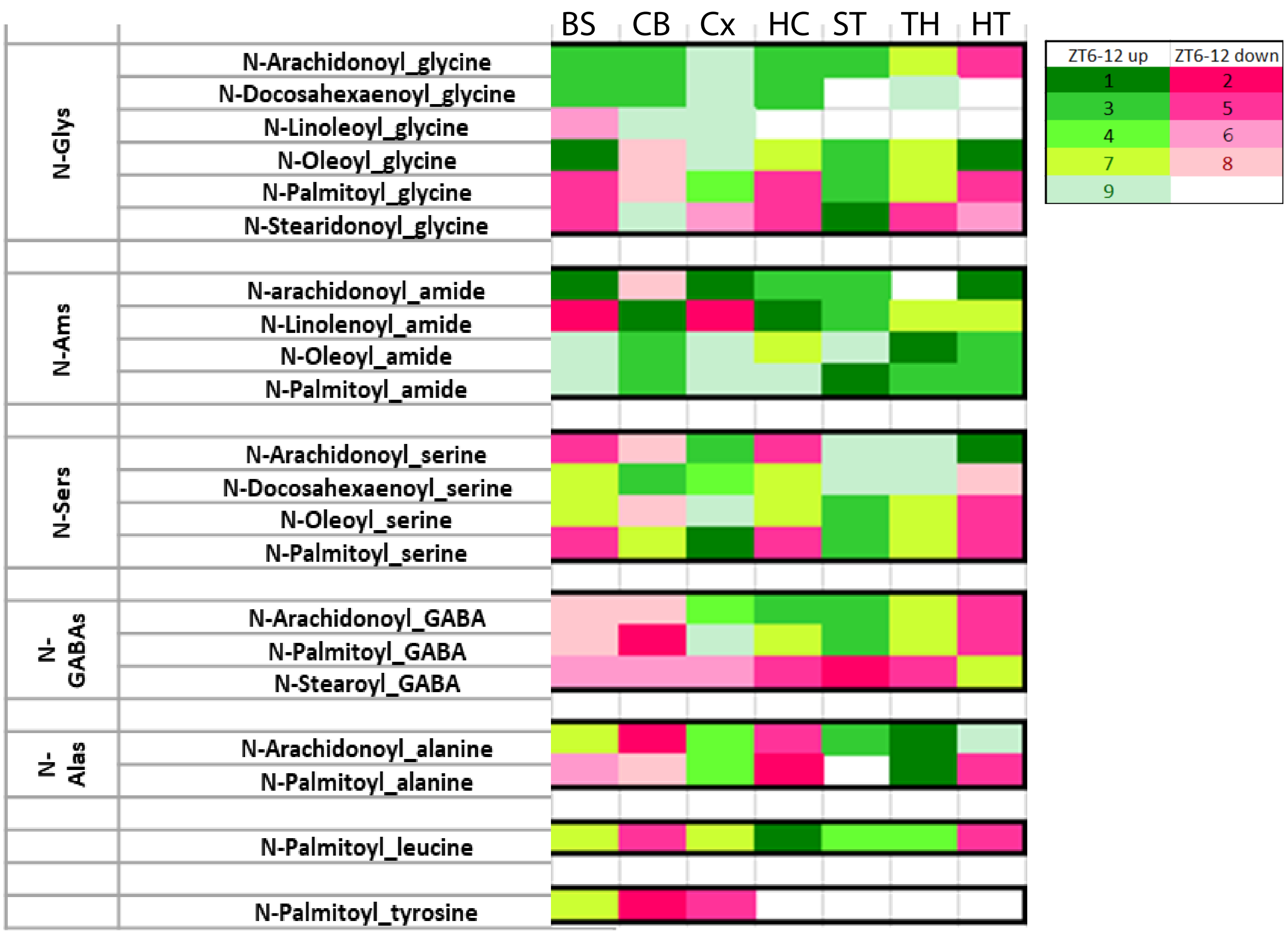
