## Supplementry figure legend for "Circadian-related Dynamics of the Endocannabinoid System in Male Mouse Brain"

**Supplementary** **Figure 1. Averaged normalized eCBs in brain regions.** **A.** Total averaged and normalized eCBs in each brain region at ZT0. *p<0.0001*, (ANOVA, Kruskal-Wallis test). **B.** FA is not significant *p=0.26* (ANOVA, Kruskal-Wallis test), **C.** 2-MAGs: *p=0.003* (ANOVA, Kruskal-Wallis test), **D.** N-Am: *p<0.0001* (ANOVA posthoc – Bonferroni), **E.** NEAs: p=0.0003 (ANOVA posthoc – Bonferroni), **F.** N-Gly*: p=0.0005* (ANOVA, Kruskal-Wallis test), **G.** N-Ser *p=0.0004* (ANOVA, Kruskal-Wallis test), **H.** N-GABA*: p<0.0001* (ANOVA posthoc – Bonferroni).

**Supplementary** **Figure 2. A-F. Individual eCBs concentration in brain regions.** Raw data is presented in the extended data table 1.

**Supplementary Figure 3.** Left: Heat-maps of total normalized averaged eCB in each brain region. Right: Quantifications of total normalized averaged eCBs in 4 transition phases in each brain region. **A.** BS (p<0.0001), **B.** Cx (p<0.0001, ANOVA, Kruskal-Wallis test), **C.** HC (p=0.176), **D.** TH (p<0.0001), **E.** HT (p<0.006). All ANOVA, Kruskal-Wallis test.

**Supplementary Figure 4:** **Averaged normalized concentrations of eCBs in different brain areas throughout the circadian rhythm. A.** Brainstem (blue), **B**. cerebellum (red), **C**. cortex (green), **D**. hippocampus (purple), **E**. striatum (orange), **F**. thalamus (black), **G**. hypothalamus (brown). Dotted colums are identified by: Triangle – FA, circle – 2-MAGs, square - NAEs, stars– N-amides, cross – N-serines, diamond – N-GABA.

**Supplementary Figure 5. Expression clusters during the circadian period based on light-to-dark transition phase.** **A.** Seven clusters showing different changes in patterns of eCBs based on whether they increase or decrease during the mid-light transition phase. Each pattern is colored differently. **B.** Cluster homogeneity scores showing how many eCBs (size) follow the same pattern (cluster). **C.** Left: eCBs differentiated by brain regions and families relative to colors of clusters in A. Right: panel showing each cluster (1-9) if increased (green-shaded) or decreased (pink-shaded) linked to colors of clusters in A. n=8/brain region.

**Supplementary Figure 6.** eCBs passed Cosinor test in brain regions. Colors of families are indicated in right bottom panel.

**Supplementary Figure 7.** **eCBs expression within sleep initiation and WEx in brain regions. A.** NEAs in the Cx. (DHEA: *p=0.0093,* ANOVA one-way, OAO: *p=0.0038*, ANOVA one-way, PEA *p=0.0045*, ANOVA one way, SEA *p=0.0121*, ANOVA Kruskal-Wallis, LEA p=0.0074 ANOVA one-way, DtEA *p=0.0021,* ANOVA one way). **B.** 2/1-DHG (*p<0.0001*, ANOVA one-way), and 2/1-EPG (*p=0.0078,* ANOVA one-way) in the CB. **C.** FAAH gene expression in the BS p=0.0024, and in the CB p=0.012; ANOVA one-way**. D.** Receptors in brain areas. Results are in supplementary materials.

**Supplementary Figure 8. eCBs expression durion ZT0, ZT1 and WEx.** Shaded colors indicate the family name as indicated in the right panel. The black square indicates the sleep effect, red square indicates the circadian effect.
